# Neural Encoding Geometry of Human Voluntary Visual Selection

**DOI:** 10.64898/2026.11.09.765268

**Authors:** Damian Koevoet, Klara Žinkovič, Stefan Van der Stigchel, Christoph Strauch

## Abstract

Selecting relevant targets from competing alternatives is a core cognitive function. While human visual selection can be tracked with high spatiotemporal precision, it remains unsolved how selected targets and non-selected alternatives are neurally encoded to effectively separate them. Building on recent work revealing the neural encoding geometry underlying target selection in animals, we here examined how the representational format of selected targets and non-selected alternatives evolve over time during human voluntary selection. Initially, selected and non-selected objects showed equal spatially selective neural activity, yet the selected target was represented more strongly after *∼*300-350 ms. Using cross-decoding, we found that selected and non-selected objects are encoded along a shared sensory axis, whereas selected targets are encoded along an additional spatially selective axis. This encoding geometry effectively separates selected targets from non-selected alternatives in neural space. Our results generalized across selection conditions: we found comparable results in instructed and free-choice selection, as well as in delayed oculomotor and covert visual selection. Together, we demonstrate that the human brain selectively encodes targets, but not distractors, along an additional axis, revealing a neural mechanism through which voluntary selection prioritizes behaviorally relevant information.

## Introduction

Environments typically contain much more content than what the visual system can process at once. This limitation is overcome by iteratively selecting relevant objects by moving gaze or shifting spatial attention to enhance the perceptual fidelity of selected content, while processing non-selected content only coarsely [1–6]. While a large body of neural and psychophysical work has uncovered the time course and perceptual consequences of human visual selection [7–16], it remains unsolved how selected targets and non-selected alternatives are neurally separated [17, 18].

Multivariate decoding techniques have enabled the tracking of visual selection with high spatiotemporal precision [19–22]. Leveraging this, studies have provided neural estimations of the time course of selection, assessed the breadth of the selected region, and tracked the fidelity of selected and non-selected targets [7, 23–26]. Moreover, visual selection has also been shown to speed up, or enhance, neural responses to stimuli presented in attended regions [13, 27, 28]. Yet, for selection to effectively guide perception and behavior, the brain must not only represent selected content more precisely, but also distinguish it from competing, non-selected information.

Extant work put forth (at least) three neural encoding geometries that could effectively separate selected targets from non-selected alternatives during visual selection. First, a straightforward neural encoding geometry is to encode selected targets and non-selected alternatives along a common sensory axis. In this case, selected targets would be disambiguated from non-selected alternatives by enhancing the sensory fidelity of only the selected target. Support for such an encoding geometry comes from recent fMRI work that concurrently decoded the position of high and low priority stimuli in a working memory task [29]. Selected targets were represented more strongly than the non-selected alternative, suggesting that differences in spatial selectivity within the same sensory axis may dissociate selected targets from non-selected alternatives. Second, recent animal experiments point to a more complex encoding geometry that recruits an additional ‘selection axis’. For example, Barbosa *et al.* [17] showed that rat primary auditory cortex (A1) encodes relevant and irrelevant features (i.e., pitch and position) along a common sensory axis during a context-based go-nogo task. Crucially, relevant, but not irrelevant, features were encoded along an additional ‘selection axis’. A similar encoding geometry has been reported in ferret A1 in a similar task [30], as well as in macaque prefrontal cortex in a paired-association task [31]. This encoding geometry seems ftexible: while the sensory axis allows to extract features from objects regardless of whether or not they are currently relevant, a selection axis allows to adaptively control the relative priority of each object based on the current task demands. Third, an opposing encoding geometry, which we term ‘deselection’, is also feasible, especially given recent proposals suggesting that non-targets can be actively suppressed in a top-down manner [32–34]. In this case, in addition to encoding selected and non-selected options onto a common sensory axis, non-selected alternatives may be actively suppressed by projecting them into a deselection axis.

Here, we aimed to characterize the encoding geometry of human voluntary visual selection. To this end, we recorded EEG and tracked selection across space and time using multivariate decoding techniques [7, 19, 21, 22, 24, 35, 36]. To uncover how the encoding geometry evolved, we concurrently tracked the position of a selected (target) and non-selected (distractor) object over time [26, 29, 37]. Through concurrent tracking of the target and distractor, we were able to arbitrate between different possible encoding geometries underlying visual selection. To ensure our findings were generalizable [38], we studied multiple selection conditions including instructed and free-choice, allowing us to chart the encoding geometry of voluntary visual selection across contexts. Studying encoding geometries, as opposed to univariate responses, should also allow for more feasible generalization of selection principles between humans and other animals.

## Results

In Experiment 1, twenty human observers completed a delayed oculomotor task (Figure 1A) while scalp EEG was recorded. On each trial, two shapes were presented in one of twenty-four possible locations around the visual field (Figure 1B). In alternating blocks, observers were either instructed to saccade to a specific shape (i.e. diamond or square) or saccaded freely to one of two shapes after a brief delay period. This allowed us to investigate instructed visual selection as typically studied in experiments, but also generalize this to more natural, free-choice selection.

**Figure 1:**
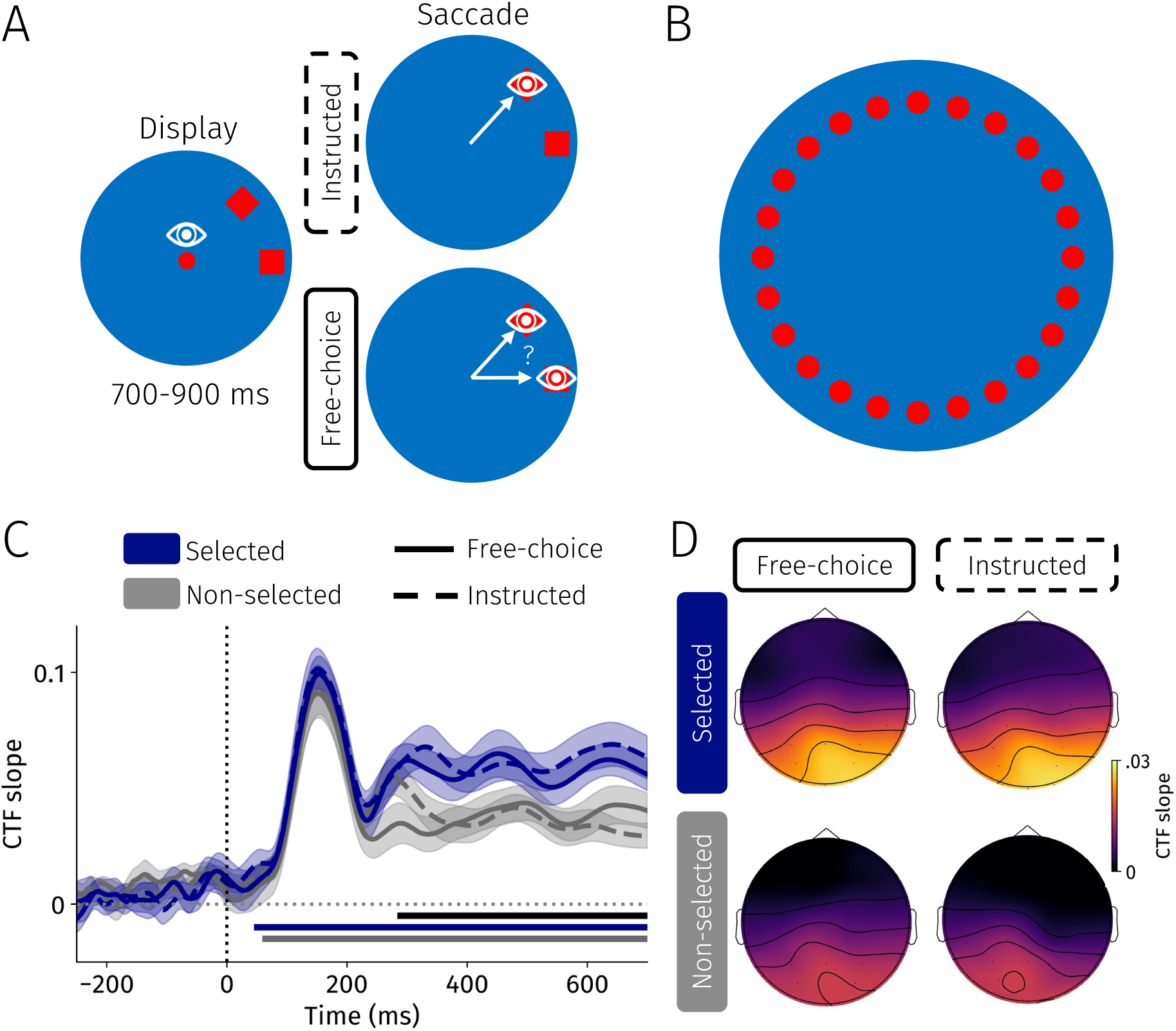
Delayed oculomotor selection task and inverted encoding modeling results from Experiment 1. **A** Task design. In instructed trials (dashed), observers selected an instructed shape; the diamond in this example. In free-choice trials (solid), observers freely selected one of the shapes. After a variable delay period, the fixation dot disappeared and observers saccaded towards the selected shape as fast as possible. **B** All possible shape locations. **C** CTF slopes (i.e. spatial selectivity) across time for selected (blue) and non-selected (gray) shapes separately for free-choice (solid) and instructed (dashed) trials. Blue and gray horizontal lines indicate significant clusters where CTF slopes were reliably above 0 (*p* < .0001) for selected and non-selected shapes, respectively. Black horizontal line indicates a significant difference between CTF slopes for selected and non-selected shapes (*p* < .001). Error bands reflect standard error of the mean. **D** Searchlight-based topographies of selected and non-selected shapes for the free-choice and instructed blocks separately. CTF slopes were averaged 300-700 ms after display onset.

Observers only rarely saccaded to the incorrect target in instructed trials (*M* = 2.3%, *SE* = 0.2%), and observers did not prefer selecting a specific shape in free-choice trials (*t*(19) = 0.04, *p* = .970, *d* = 0.01; see Supporting Figure 1 for more behavioral details). Thus, observers were were well able to perform the task.

### Time-Resolved Neural Tracking of Visual Selection

We aimed to track the position of the selected and non-selected shapes, allowing us to compare their spatial representations across time (Figure 1). To this end, we employed inverted encoding modeling (IEM) [19, 22], which has previously been used to track the allocation of spatial attention with high temporal precision [20, 24, 26, 27, 39]. We focused our neural analyses on the display period because no saccades were executed yet. This allowed us to concurrently decode the position of both shapes without ocular artifacts.

We found strong spatially selective neural activity shortly after display onset to both the selected and non-selected shapes which persisted throughout the delay period (both *p*s < .0001; Figure 1C). We were thus able to concurrently track the spatial selectivity to both shapes, and could examine how they evolved across time. We found that spatially selective neural activity tracked the position of both shapes in two phases. Spatially selective neural activity to the selected and non-selected shapes were initially comparable during sensory processing. Only later did we observe sharper selectivity for the selected target over the non-selected alternative (*p* < .001), suggesting that observers had selected targets *∼*300 ms after display onset in both instructed and free-choice trials. We found no reliable differences between instructed and free-choice selection, nor did we observe a reliable interaction (i.e., selected/non-selected x instructed/free-choice).

To further characterize this spatially selective activity, we visualized topographies from the IEM analyses using a searchlight approach [Figure 1D; 40]. Specifically, we iteratively selected groups of 5 neighbouring electrodes and reconducted the IEM procedure for each group separately [24]. To simplify visualizations, we averaged reconstructed CTF slopes 300-700 ms after display onset. We computed the average CTF slope (i.e., spatial selectivity) for each electrode by averaging across all electrode groups that a specific electrode was included in. This allowed us to visualize spatial selectivity collapsed across all positions for each electrode separately. In line with prior work, we found that spatial representations of selected and nonselected shapes were both predominantly represented in posterior electrodes [24, 25], and topographies were similar during instructed and free-choice selection.

Neural decoding effects can, in some cases, be caused by miniature gaze patterns instead of neural activity [41]. This may be especially problematic when decoding spatial contents, as gaze can point toward where attention is allocated [42, 43]. We therefore conducted control analyses and ruled out gaze patterns as an alternative explanation for all decoding results in this paper (Supporting Figure 2).

**Figure 2:**
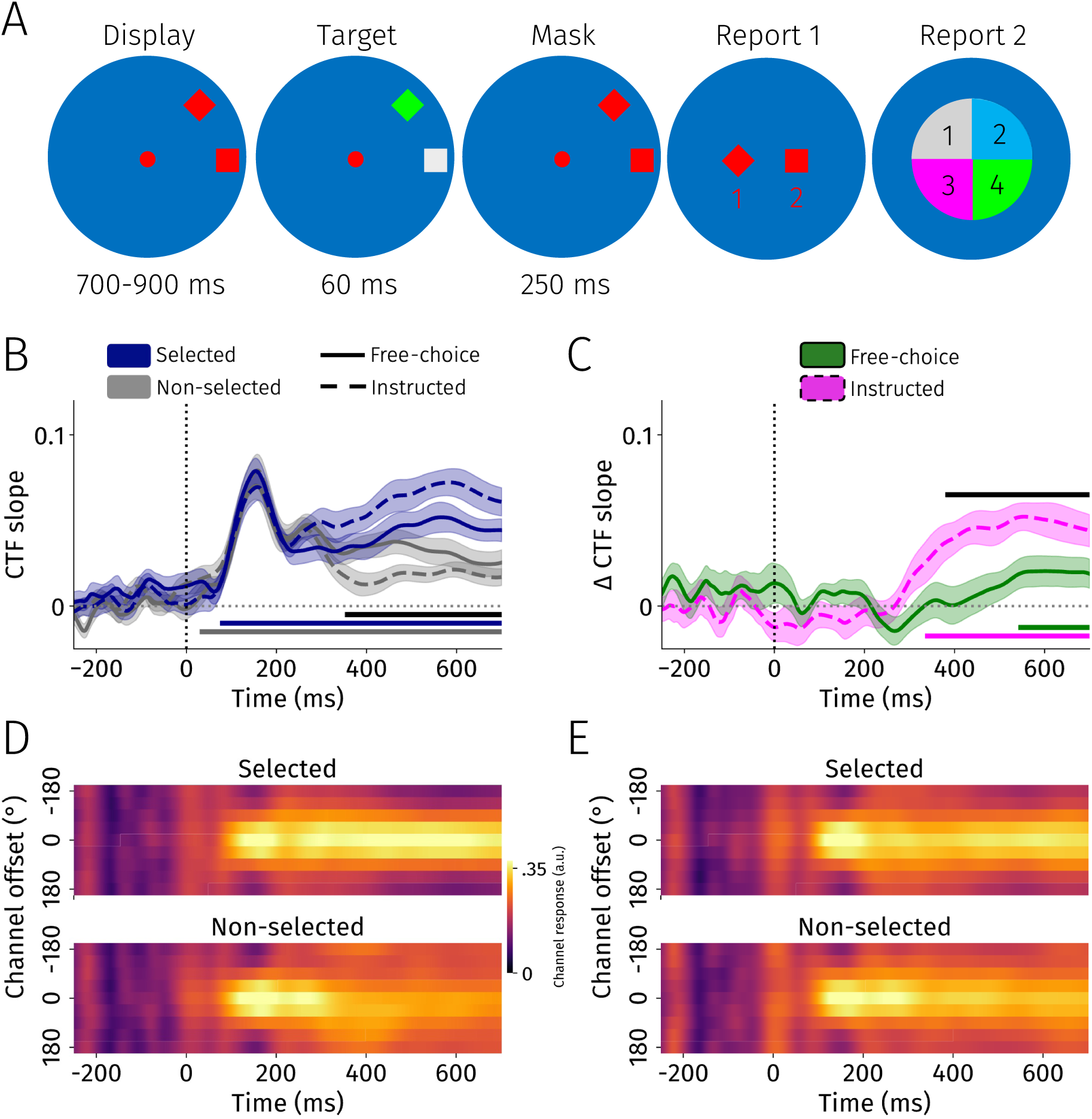
Color identification task and inverted encoding modeling results of Experiment 2. **A** Task design. In instructed trials, observers selected an instructed shape. In free-choice trials, observers freely selected one of the shapes. After a variable delay period, both shapes briefly changed color. Observers indicated the selected shape, and subsequently to which color it changed during the target period. **B** CTF slopes (i.e. spatial selectivity) across time for selected (blue) and non-selected (gray) shapes separately for free-choice (solid) and instructed (dashed) trials. Blue and gray horizontal lines indicate significant clusters where CTF slopes were reliably above 0 (*p* < .0001) for selected and non-selected shapes, respectively. Black horizontal line indicates a significant difference between CTF slopes for selected and non-selected shapes (*p* < .0001). Error bands reflect standard error of the mean. **C** Difference traces for the free-choice (green) and instructed (purple) conditions. Traces were calculated by subtracting non-selected traces from selected traces Positive values indicate higher CTF slopes for selected than non-selected shapes (and vice versa). Black horizontal line indicates a significant difference between difference traces (i.e., interaction; *p* < .01). **D** Heatmap of modeled channel responses for each position bin for the instructed condition. Channel offset of 0° indicates the relevant position, and higher absolute channel values indicate a larger distance from that position. **E** Same as D but for the free-choice condition. All error bands reflect standard error of the mean.

Together, these data characterize the temporal dynamics of visual selection. Initially, both eventual targets and alternatives are represented at equal strength during sensory processing, and only after selection has occurred is the selected object represented more sharply than the non-selected alternative.

### Tracking Visual, Not Motor, Selection

In Experiment 1, we were able to track how voluntary visual selection evolved over time by concurrently tracking the position of two shapes. However, competition between objects can occur at the visual or motor stage [44, 45]. Since the position of a shape always corresponded with the direction of a saccade, it remains unclear whether the above results can be attributed to visual or motor selection (or a combination of both). Compounding this possible issue, we recently found that that oculomotor-related spatially selective activity is captured by the EEG-decoding techniques used here [24]. We circumvented this possible limitation in Experiment 2 by decoupling visual selection from motor preparation.

Twenty-one observers performed a color identification task wherein they selected an instructed shape or freely selected one of two shapes in alternating blocks (Figure 2A). Observers monitored the selected shape for a change in color, which was briefly masked afterwards. In contrast to Experiment 1, observers used a manual key response to indicate whether they selected the square or diamond stimulus, effectively separating visual selection from motor preparation. Observers reported the color of the selected shape to conclude the trial.

All observers identified target colors above chance level (25%) in both instructed (*M* = 75.5%, *SE* = 1.7%) and free-choice blocks (*M* = 79.0%, *SE* = 1.8%; Figure 2D). Accuracy was higher in free-choice compared with instructed blocks (*t*(20) = 3.86, *p* < .001, *d* = 0.84). As in Experiment 1, observers selected the wrong shape only rarely in instructed trials (*M* = 3.3%, *SE* = 0.7%), and no significant preference for a specific shape was observed in free-choice trials (*t*(20) = 1.61, *p* = .123, *d* = 0.35; see Supporting Figure 1 for more details). Thus, observers performed the task well.

Turning to the neural data (Figure 2B-E), we could reliably track the spatial position of both the selected and non-selected shapes (*p*s < .0001). As in Experiment 1, we again found two phases of spatially selective neural activity. While the selected and non-selected shapes were represented at similar strengths shortly after display onset, the selected shape was represented more strongly than the non-selected shape later in the trial (*∼*350 ms after display onset; *p* < .0001; Figure 2B).

We also observed a reliable interaction effect: the difference in CTF slope between selected and non-selected was larger in instructed compared with free-choice trials (*p* = .008; 376-700 ms; Figure 2B-E). Nonetheless, we still observed sharper spatially selective activity to the selected compared with the non-selected shape in the free-choice condition. Indeed, selection seemed to occur later in free-choice than in instructed trials, possibly because observers spent longer to decide which shape to select. There was no significant difference in CTF slope between instructed and free-choice trials, implying that the ‘total amount’ of spatial selectivity was similar between conditions.

Together, these data extend the findings from the oculomotor domain to covert visual selection and effectively rule out motor preparation as a compelling alternative explanation to our results.

### Encoding Geometry of Visual Selection

Until now, we have demonstrated that neural decoding tracks voluntary visual selection and that it follows a two-phase process. But *how* does selection separate the selected from the non-selected shape? We considered the three aforementioned encoding geometries (Figure 3A) [based on 17].

**Figure 3:**
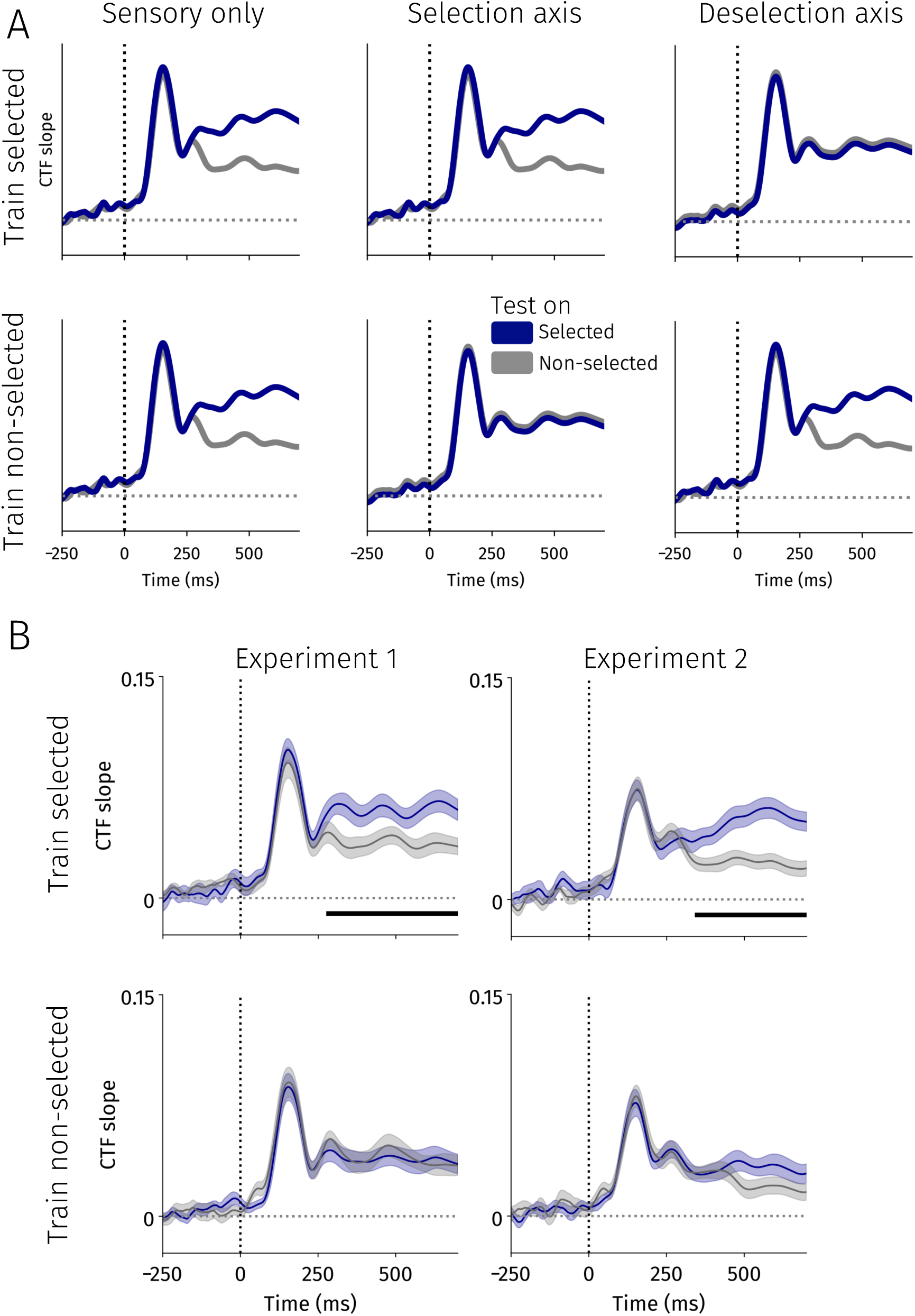
Arbitrating between possible encoding geometries. **A** Hypothetical encoding geometries of voluntary visual selection. Predicted CTF slopes (i.e. spatial selectivity) across time split based on the training and testing conditions for sensory only, selection, and deselection accounts. **B** Cross-decoding results from Experiments 1 (left) and 2 (right). CTF slopes across time split per training condition, and split based on whether we tested the model on selected (blue) or non-selected (gray) positions. Horizontal black lines indicate significant clusters between selected and non-selected CTF slopes (*p*s < .001) Error bands reflect standard errors of the mean.

To arbitrate between these encoding geometries, we adopted a cross-decoding approach [17, 24]. Specifically, we either trained decoders (IEMs) using only the position of the selected (as in all previous analyses; Methods) or the position of the non-selected shape. We hypothesized that a sensory-only account would result in a stronger spatial representation of the selected target when training on either the selected or non-selected shape’s position. By contrast, a selection-axis account predicts that when decoders are trained on the selected shape’s position but not on the non-selected shape’s position, the selected shape should be represented more strongly than the non-selected shape. A deselection-axis account predicts the inverse: the non-selected shape should be represented more weakly than the selected shape, but only when the decoders are trained on the position of the nonselected shape. Note that since we found no differences between the instructed and free-choice blocks in the following analyses, we collapsed across these conditions to increase power.

We observed spatial selectivity to the selected and non-selected shapes (all *p*s < .001) in all conditions (Figure 3B,C). This demonstrates that a common sensory axis codes for stimulus position regardless of whether a shape was selected (as predicted by all encoding geometries under consideration here). Disambiguating between encoding geometries, we observed that selected targets were more strongly represented than non-selected targets when the decoders were trained on the position of the selected target (*p*s < .001). In contrast, we found no reliable difference in spatial representation between the selected and non-selected targets when decoders were trained on the position of the non-selected target. We directly compared the difference in spatial selectivity between selected and non-selected targets between training conditions (i.e., interaction) and found a consistent pattern across experiments (Exp. 1: *p* = .002; Exp. 2: *p* = .048).

These results support a selection-axis account: selected and non-selected objects are encoded along a common sensory axis, yet selected targets (but not nonselected alternatives) are encoded onto an additional selection axis. But does this encoding geometry effectively separate selected targets from non-selected alternatives in neural space? To investigate this, we examined the neural geometry of the spatially selective neural activity by projecting reconstructed CTFs during an early (0-200 ms) and late (500-700 ms) time window into a two-dimensional principle component analysis (PCA) space (Figure 4) [46, 47]. The projected data closely resembled the circular structure of our task for both selected and non-selected shapes in early and late time windows (Figure 4A). However, we observed that the circular structures of selected and non-selected shapes were distant in neural space during the late but not during the early window, and only when trained on the selected position (Figure 4A, bottom left).

**Figure 4:**
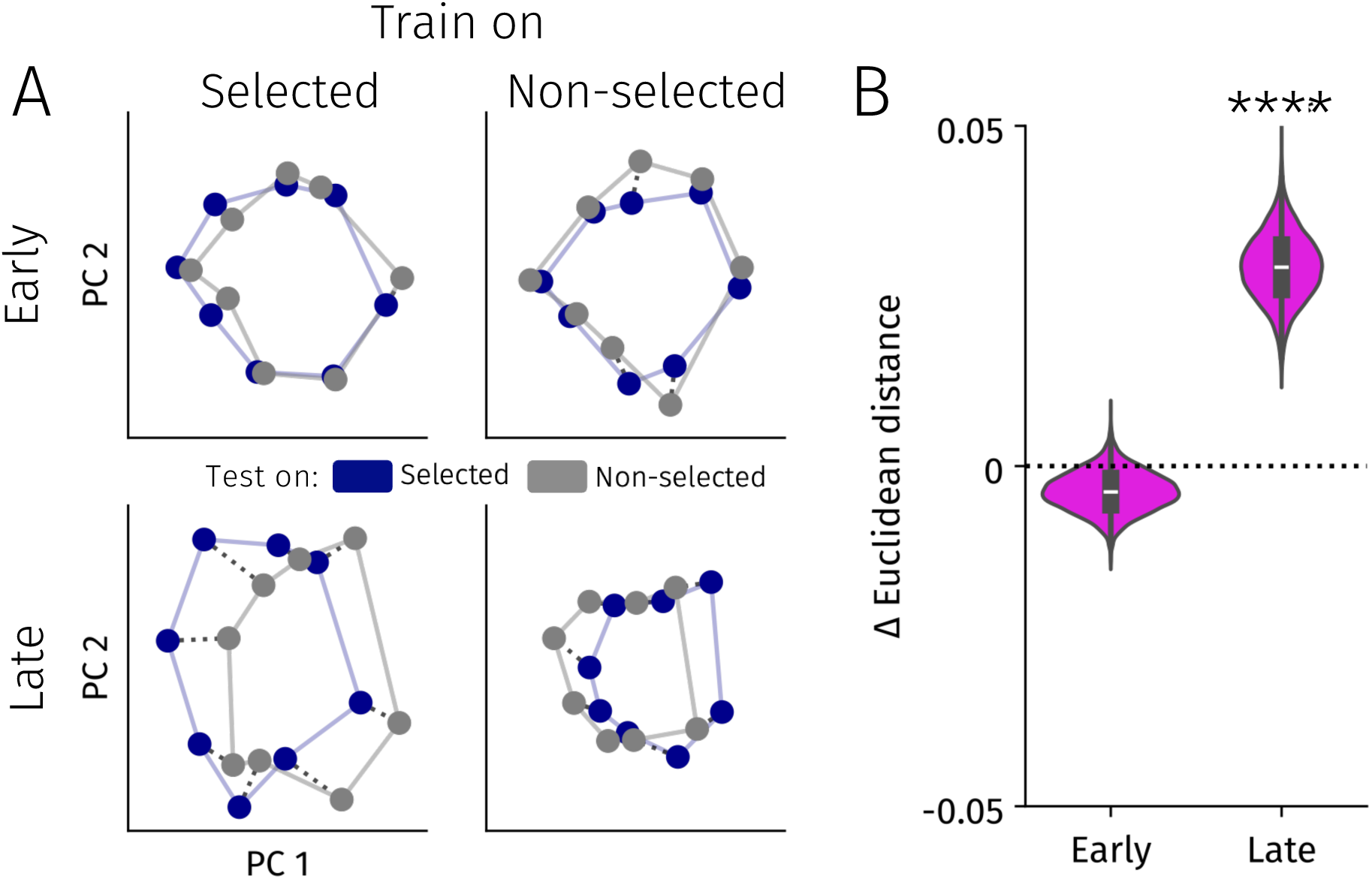
Neural geometry of voluntary visual selection. **A** Reconstructed CTFs projected into a two-dimensional PCA space to visualize the neural geometry of spatial selectivity. Reconstructed CTFs were projected to PCA space in an early (0-200 ms) and late (500-700 ms) time window separately per training condition. Selected and non-selected targets are represented in blue and gray, respectively. Thick lines indicate interconnected positions, which reveal a circular pattern of the data, resembling the positioning of stimuli in the tasks. Dash lines indicate distances between the same positions based on whether it was selected or non-selected. **B** Euclidean distance between the same positions based on whether it was selected or not was calculated separately per training condition (selected or non-selected) and time window. Positive values reflect greater difference (i.e. bigger shifts in neural geometry) when training on the selected compared with the non-selected positions. Violin distribution reflects the 10,000 train-test splits used for the permutation test. \*\*\*\**p* < .0001.

To formally test this, we calculated the Euclidean distance in PCA space for each position between when this position was the selected or non-selected position. If recruiting an additional selection-axis facilitates the separation of selected and non-selected spatial representations, Euclidean distances in PCA space should be larger between selected and non-selected positions when decoders were trained on selected positions. We tested this using a cross-validated permutation test (10,000 train-test splits, Methods). In the early time window, we found no significant difference in Euclidean distance between selected and non-selected shapes between training conditionsx (*p* = .905; Δ*_Euclidean_*=−.003, *SD* = .003; Figure 4B). In the later time window, when selection had taken place, we found a larger distance in PCA space between selected and non-selected shapes when decoders were trained on selected compared with non-selected positions (*p* < .0001; Δ*_Euclidean_*=.029, *SD* = .0049; Figure 4B). This demonstrates that recruiting an additional selection axis facilitates the separation of selected targets from non-selected alternatives in neural space.

## Discussion

We here investigated the underlying encoding geometry through which the human brain parses visual input into selected targets and non-selected alternatives during visual selection. Specifically, we applied multivariate decoding techniques to EEG data to track selected and non-selected representations to uncover when and how visual selection evolved across time. This allowed us to characterize the temporal dynamics and probe the neural encoding geometry of human voluntary visual selection. To ensure our findings were generalizable [38], we investigated multiple selection contexts, including instructed and free-choice, as well as oculomotor and covert spatial selection.

Our main goal was to uncover the neural encoding geometry of human voluntary visual selection. Using cross-decoding [17, 24], we found that selected and nonselected shapes were encoded along a shared sensory axis, yet selected targets were encoded onto another spatially selective ‘selection’ axis. This suggests that voluntary visual selection is primarily achieved by boosting the representation of the selected target, not through suppression of the non-selected alternative. Further analysis of the spatially selective activity revealed that encoding a selected target onto an additional axis facilitates the separation of selected targets from competing alternatives. This encoding geometry may allow for more efficient action selection [see 48]. Given the large differences across tasks, it is remarkable that comparable encoding geometries of target selection have been reported in rats, ferrets, non-human primates [17, 30, 31], and now in humans. The observation of similar encoding geometries across species and tasks suggests this may provide a useful framework for identifying general principles of target selection, even when the specific neural responses differ [49–52].

We found that visual input is parsed in two stages: an initial sensory processing phase where both stimuli were represented at similar strengths, followed by a selection phase *∼*300-350 ms after display onset where the selected target was represented more strongly than the non-selected alternative. This temporal profile dovetails with previous behavioral and neural estimates of the time course of voluntary selection [7, 9–12]. Moreover, the two stages reported here are compatible with dynamic theories of visual attention where distinct factors affect selection at different timescales [53]. Indeed, mounting evidence suggests that physical salience attracts attention rapidly, while voluntary goal-driven control steers selection later [54–57]. The currently employed approach of concurrently tracking two potential selection targets may enable to uncover how stimulus-, history-, cost-, and voluntary goal-based factors dynamically shape attentional priority over time [57–59].

The current data also add to a growing body of work investigating free-choice visual selection [sometimes referred to as ‘willed attention’; 60]. We extend this here. First, while prior work was able to discern between whether attention was allocated to the left or right visual hemifield [61–63], we here tracked the precise position of objects around the visual field. Second, we here tracked both selected and nonselected objects which allowed us to study visual selection while controlling for sensory input to obtain a more accurate temporal profile of selection. Extending instructed voluntary selection to free-choice selection bridges our findings closer to application in the form of brain-computer interfaces, as well as to more natural viewing behavior.

We found that visual selection unfolded similarly in instructed and free-choice selection in Experiment 1. Yet, we observed a stronger representation of the selected shape compared with the non-selected object in instructed compared with free-choice selection in Experiment 2 (Figure 2C). Although these results from Experiment 2 align with a previous study [62], we are uncertain what drove the difference between experiments. One possibility is that free-choice visual selection may be resolved more quickly if competition can (also) be resolved at the motor level [e.g., 45]. Another option is that observers delayed their decision until later in the trial in Experiment 2, plausibly consistent with the higher accuracy in free-choice compared with instructed trials.

We note that, in some cases, cross-decoding asymmetries as those reported here (Figure 3) may reflect differences in signal-to-noise ratio between conditions instead of revealing ‘real’ neurocognitive processes [64]. Our results are unlikely to be explained by this concern because 1) selected and non-selected shapes were presented simultaneously in the same trial, and 2) because selected and non-selected shapes were represented at similar strengths briefly after display onset indicating similar signal-to-noise ratios. We therefore conclude that the reported cross-decoding asymmetry reflects a true property of the neural activity during visual selection, not a signal-to-noise ratio artifact.

How does the brain effectively parse visual input into relevant, selected targets and competing, non-selected targets? Across selection contexts, relevant and irrelevant inputs are initially processed at equal strength, but subsequently selected targets are represented at a higher fidelity than non-selected alternatives. To achieve this boosted representation, selected targets, but not non-selected alternatives, are encoded along an additional spatially selective selection axis. This effectively separates relevant visual contents from competing alternatives in neural space, which may facilitate the transformation of visual representations into sensible actions.

## Methods

Data and analysis code is available at: https://figshare.com/s/881cd99093ab1eec353b.

### Observers

We set our target sample to 20 based on prior work concurrently tracking two positions using multivariate EEG decoding techniques [26]. This number is also comparable or exceeds past work using multivariate decoding to track spatial attention [7, 20, 24]. For each experiment, we collected data from twenty-five observers. In Experiment 1, we discarded data from three observers due to technical issues (i.e., data were not stored or experiment crashed), from one observer because they did not finish the full experiment, and from one observer because they had less than 300 artifact-free trials in at least one of the two conditions. In Experiment 2, we discarded data from one observer due to low task accuracy and data from two other observers were discarded due to a high number of trials with artifacts (< 300 artifact-free trials for at least one condition). This left final samples of 20 (*M_age_* = 22.60, range: [19–26], 15 female, 5 male) and 21 (*M_age_* = 23.09, range: [19–29], 17 female, 4 male) observers for analyses of Experiments 1 and 2, respectively. All observers reported normal vision and no history or diagnosis of attention-deficit hyperactivity disorder or autism spectrum disorder. We slightly overshot our target sample for Experiment 2 because we performed artifact rejection in batches. Observers were compensated for participation with course credits or received €8 per hour. Experimental procedures were approved by the Utrecht University ethical review board (approval code: 24-0197).

### Procedures

#### Experiment 1

Observers performed a delayed oculomotor selection task (Figure 1A). In alternating blocks, observers were either instructed to select a specific shape (informed between blocks) or could freely select one of the two shapes.

Each trial started with a fixation period that ranged between 750-1000 ms. Observers were instructed to fixate a central red (HSV: 0.1.0.93) dot (0.5°) a blue (HSV: 240.1.1) circle that was presented on a black background. Red and blue colors were matched in luminance. Then, two shapes appeared (square or diamond, 0.8° x 0.8°) in one of twenty-four locations around the visual field at an eccentricity of 10°. Positions progressed in steps of 15° starting at 0°. The two shapes were separated by at least 30° to limit saccade averaging [65]. In instructed trials, we always presented one square and one diamond. In free-choice trials, we randomly chose shape identities to prevent observers from simply selecting a shape throughout a block. After a variable delay period of 700-900 ms, the fixation dot disappeared, signaling observers to saccade toward the selected shape as fast as possible. The experiment consisted of a total of 20 blocks, each containing 60 trials. For both experiments, we counterbalanced whether observers started with an instructed or free-choice block, and participants completed two practice blocks prior to the experimental session.

#### Experiment 2

The procedure for Experiment 2 was closely based on Experiment 1. Observers completed a color identification task (Figure 2A). In contrast to Experiment 1, we equalized visual input by always presenting one square and one diamond in both instructed and free-choice trials. Moreover, as we wanted to ensure that (oculo)motor preparation could not account for our results, observers were instructed to maintain fixation throughout each trial. We aborted trials when gaze deviated more than 3° from the central fixation point.

After the display period, both shapes briefty changed to one of four possible target colors for 60 ms (lightgreen, magenta, gray, or cyan). Target colors of the two shapes were never the same. Following a brief mask display (250 ms), observers first reported which shape they selected. We randomized the mapping between keys (1 or 2) and shape on each trial to prevent any effects of motor preparation during the display period. Then, observers reported which color the selected shape had turned into.

The experiment consisted of a total of 18 blocks, each containing 60 trials. We slightly reduced the number of trials because they were slightly longer than those in Experiment 1.

### Data Acquisition and Processing

Gaze and pupil data were recorded from the right eye at 1000 Hz with an EyeLink 1000+ (SR Research, Mississauga, Ontario, Canada). Observers were positioned in a chinrest 67.5 cm from the monitor (ASUS ROG PG278Q; 2560 x 1440; 100 Hz). The eye-tracker was calibrated using a 9-point procedure, and recalibration was performed whenever necessary between blocks. Stimuli were presented using PsychoPy (v2024.2.4) [66].

EEG was recorded from 30 active electrodes (2,048 Hz) using a BioSemi ActiveTwo system (BioSemi, Amsterdam, The Netherlands). Note that EEG data was recorded at 4,096 Hz for some observers in Experiment 2. However, because data were downsampled to 512 Hz at the beginning of the analysis pipeline, this did not affect results. We recorded from International 10/20 sites: Fp1, AF3, F7, F3, FC1, FC5, C3, CP1, CP5, P7, P3, Pz, PO3, O1, Oz, O2, PO4, P4, P8, CP6, CP2, C4, FC6, FC2, F4, F8, AF4, Fp2, Fz, and Cz. Additionally, electrodes were placed *∼*1 cm from the outer canthi of the left and right eyes to record horizontal EOG. We also placed electrodes above and below the right eye to monitor vertical EOG. Electrodes were placed on the left and right mastoid bones that were used for later offiine re-referencing.

#### Glitches

Due to experimenter error and technical issues, a limited number of trials at the start of the experiment were not recorded in data from three observers, and for two observers either one or two full blocks were not recorded in the data in Experiment 1. In Experiment 2, we lost data from two blocks for one observer due to technical issues. Nevertheless, all included observers had at least 300 artifact-free trials in both the instructed and free-choice blocks.

#### Data Processing and Artifact Detection

Data processing and analyses were performed using custom Python code, and were closely based on previous work [24, 59]. We first re-referenced EEG data to the average of the left and right mastoids and removed line noise and its harmonics using a Notch filter. After downsampling the EEG data to 512 Hz, data were highpass filtered at 0.05 Hz to remove slow drifts from the data. Data were then epoched −250 −700 ms around display onset.

We detected artifacts with an automated detection algorithm from the *autoreject* Python package [67]. Briefty, instead of relying on predefined peak-to-peak thresholds to reject data, the detection algorithm uses cross-validation to estimate peak-to-peak thresholds on a by-electrode basis per observer. Based on these personalized peak-to-peak thresholds, the algorithm detects trials containing artifacts. Whenever possible, bad electrodes within a specific epoch were corrected using spherical spline interpolation [68].

In addition, we detected ocular artifacts using eye-tracking data [based on 24, 27]. First, any trial with missing pupil size data was discarded to remove blink trials. Second, we calculated drift-corrected gaze data by subtracting the median gaze position prior to display onset (−250 - 0 ms). We discarded trials where drift-corrected gaze deviated more than 1° from central fixation [24, 69]. We also computed a sliding window of 60 ms that progressed in steps of 10 ms. If the difference in mean gaze position in the first and last halves of the sliding window exceeded 0.5°, the trial was discarded. Lastly, we visually inspected EEG data for all channels to confirm data quality. The artifact rejection process was effective as remaining deviations from central fixation were minimal (<0.10°; Supporting Figure 2).

After further exclusion of trials where observers selected the wrong shape in instructed trials, an average of 945.50 (*SD* = 123.78) and 792.52 (*SD* = 116.57) trials remained for analyses in Experiments 1 and 2, respectively.

Prior to our decoding analyses, we baseline corrected the voltage data by subtracting the mean voltage activity −250-0 ms prior to display onset. In keeping with past decoding work using voltage, we applied a 6 Hz lowpass filter to increase signal-to-noise ratio [59, 70]. To ensure data did not shift across time, we employed a zero-phase FIR filter, which applies the filter both forward and backward to eliminate phase shift (using *filter_data* from the *mne* package). These baseline-corrected, filtered data served as input to our decoding analyses.

### Inverted Encoding Models

We aimed to concurrently track the position of selected and non-selected shapes using inverted encoding models [IEM; 19, 22]. We chose IEMs as these models have been demonstrated to closely track spatial attention at a high temporal resolution [7, 20, 24, 71, 72]. Moreover, as IEMs have previously been employed to track two positions concurrently [26, 59, 73], they were well-qualified for the current purpose. We wanted to track the position of both the selected and non-selected shapes because this allowed us to study when and how visual selection unfolded across time.

Our IEM approach was based on prior work [7, 24, 58, 59, 71]. We reconstructed spatially selective channel tuning functions from the voltage topographies across the scalp. Although our tasks included twenty-four different positions, we grouped trials into equal bins consisting of three neighbouring positions to create eight position bins [as in 7, 72]. We assumed that voltage reflected the weighted average of these eight spatially sensitive channels, where each channel is spatially selective to a specific position. Basis channel tuning functions were modeled using a half-sinusoid raised to the 7th power which were each centered to one of eight position bins around the visual field.

We split the data into one training and one test set in each iteration. For each set, we ensured an equal number of trials within the eight selected shape position bins, as well as an equal number of trials from the instructed and free-choice conditions. We fit the basis channel tuning functions to the neural data using a general linear model (using *numpy.linalglstsq*). In our first set of analyses, we specifically fit the encoding model using the selected shape positions during training [58]. This ensured that the encoding model was trained on the same data across all conditions, and that the only difference between conditions was the data that used for later reconstruction [22]. The procedure for the crossdecoding analysis was identical, but we then also exclusively trained on the non-selected shape’s position. We inverted the weights from the general linear model, which allowed us to reconstruct putative spatially-sensitive channel tuning functions separately for all four conditions (selected/non-selected x instructed/free-choice) of interest. For all analysis but the PCA analysis (see below), reconstructed channel tuning functions were re-centered so that the center of the function corresponds to the position of interest. In order to consider all trials in this procedure, we ran 100 iterations. To obtain time-resolved measures of spatial selectivity, we repeated this procedure for every sample (every *∼*1.96 ms).

We quantified spatial selectivity by calculating the slope of reconstructed channel tuning functions. After averaging channel responses across positions equidistant to the position of interest, we used linear regression to estimate channel tuning function slopes. Thus, higher slope values indicate stronger spatially selective neural activity.

### Principle Component Analysis

We wanted to examine whether encoding the selected target along an additional ‘selection’ axis would separate the selected target from the non-selected option in neural space. We therefore took the non-centered reconstructed CTFs from the inverted encoding modeling analysis, and examined its neural geometry. We first averaged across observers (but not iterations) to increase signal-to-noise ratio and to examine group-level coding. For the visualization only, we also averaged across all iterations of the IEM decoders. A matrix of 32 conditions (8 positions x 2 training conditions x 2 testing conditions) by 8 CTF responses (i.e., channel responses for each position) was constructed. We then created the two-dimensional PCA space using singular value decomposition (as in implemented in *sklearn*), resulting in a 32 conditions by 2 PC matrix.

We tested for differences in neural geometry between training conditions by using a cross-validation permutation test approach. To this end, data were split into training and testing blocks (50%) separately for CTFs trained on the selected and non-selected shape’s position. Blocks were averaged across iterations, and concatenated resulting in a matrix of 32 conditions (8 positions x 2 training conditions x 2 testing conditions) by 8 CTF responses for the training and test blocks. To avoid train-test leakage, we fit the PCA using the training data, and projected the testing data into the resulting PCA space. Then, for each of the eight possible stimulus positions in our task, we calculated the Euclidean distance between that position in PCA space whether it was selected or not selected. We calculated these Euclidean distances separately for each training condition, and subsequently compared these distances across 10,000 train-test splits. We obtained *p*-values by counting the amount of train-test splits where the Euclidean distance between selected and non-selected shapes was larger when decoders were trained on the selected shape’s position compared with when trained on the non-selected shape’s position.

This entire procedure was performed for the averaged reconstructed CTFs in an early (0-200 ms) and late (500-700 ms) time window, allowing us to determine whether the neural geometry effects were specific to selection instead of early sensory processing.

### Statistical Analysis

For the analysis of behavioral data, we used paired-samples *t*-tests to analyze differences between the instructed and free-choice conditions. Where appropriate, we report Cohen’s *d_z_* as a measure of effect size [74].

We tested whether CTF slopes were reliably above 0 (indicating reliable spatial selectivity) across observers. We addressed issue of multiple comparisons by employing cluster-based permutation tests [75]. Following prior work [24, 59], we used one-tailed tests (using *permutation_cluster_1samp_test* from the *mne* package) to identify cluster candidates and test their significance against 0 separately for each condition. For ease of inference, we tested whether CTF slopes were reliable to the selected and non-selected shapes collapsed across instructed and free-choice trials. This did not affect our results: testing separately per each of the four conditions revealed reliable CTF slopes for all conditions in both experiments (all *p*s < .001). We also compared differences in CTF slope between conditions using cluster-based repeated-measures ANOVA permutation tests. All cluster-based tests were conducted at *α* = .05 using 10,000 iterations.

## Author contributions

Conceptualization: D.K. and C.S.; methodology: D.K., K.Z., S.V.d.S. and C.S.; software: D.K.; investigation: D.K. and K.Z.; formal analysis: D.K.; writing—original draft: D.K.; writing—review and editing: D.K., K.Z., S.V.d.S. and C.S.; funding acquisition: S.V.d.S.; resources: S.V.d.S.; supervision: S.V.d.S. and C.S.

## Acknowledgments

We thank Vicky Voet for helping with data collection. We thank Dirk Van Moorselaar for discussions about the inverted encoding modeling analysis.

## Data and code availability

Data and analysis code is available at: https://figshare.com/s/881cd99093ab1eec353b

## Supporting Information

**Supporting Figure 1:**
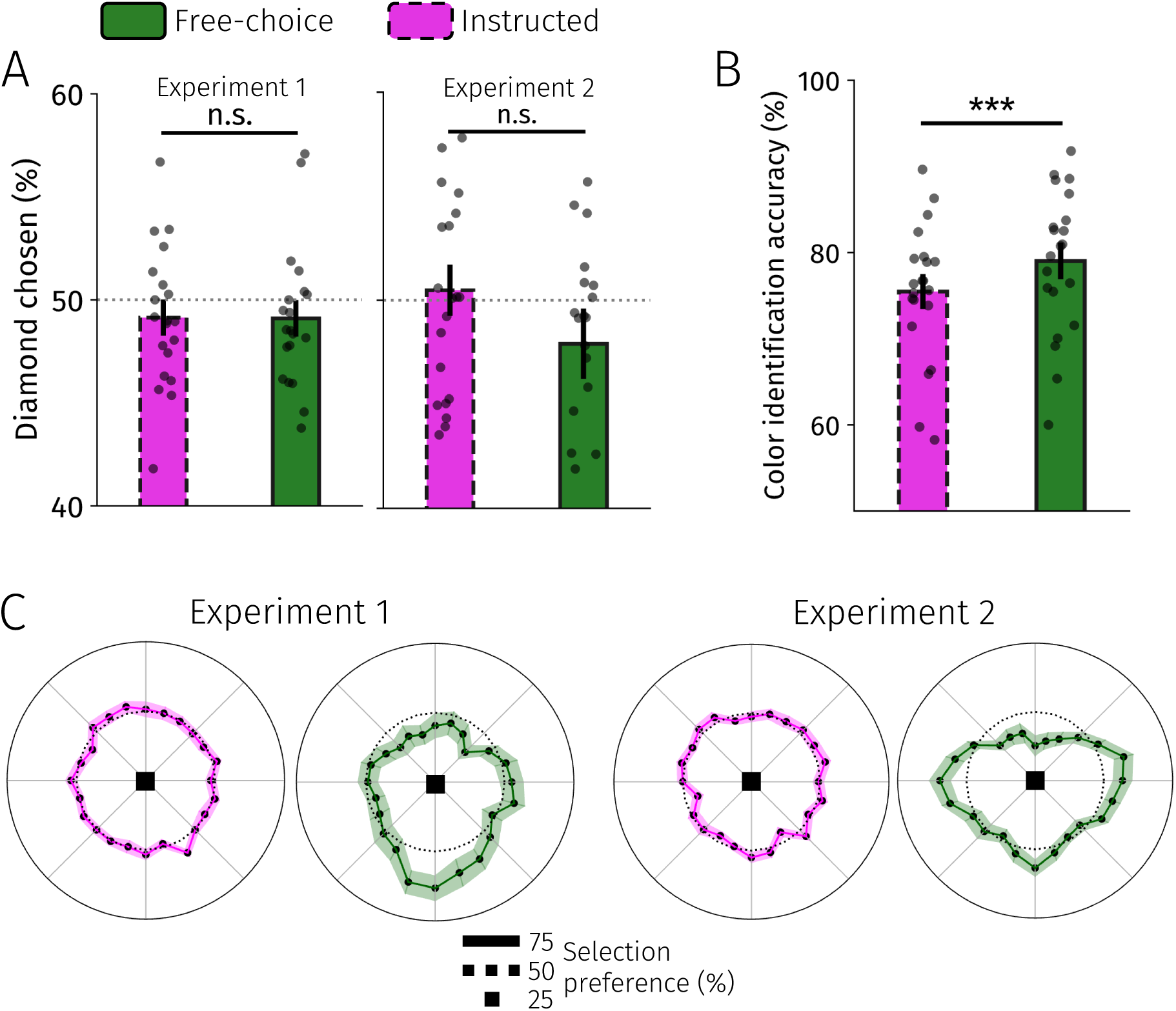
Detailed behavioral analyses. **A** Observers did not prefer selecting a specific shape (diamond vs. square) in Experiments 1 (left) and 2 (right). **B** Observers performed well above chance in the color identification task. Accuracy was higher in free-choice compared with instructed blocks. **C** Selection preferences across the visual field in both Experiments. We quantified selection preferences by calculating the percentage of trials that observers chose a specific position relative to the total times it was offered. By design, selection preferences were uniform in instructed blocks (slightly off because of trial rejections). All error bars reflect standard error of the mean.

**Supporting Figure 2:**
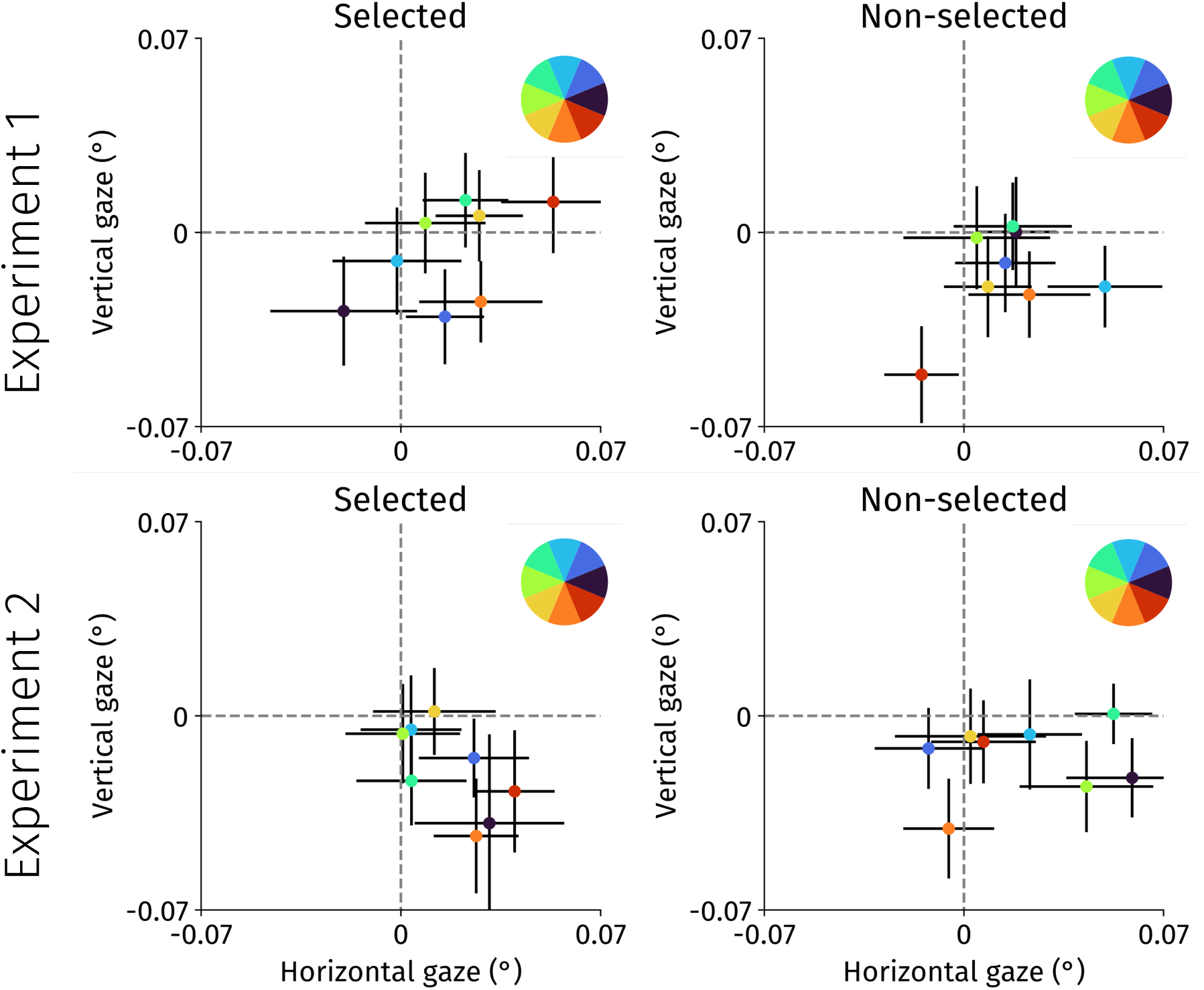
Eye-movement control analysis. To investigate whether gaze patterns affected the neural decoding results, we computed drift-corrected gaze position by subtracting the mean gaze position (−250-0 ms before display onset), and plotted the averaged (200-700 ms) gaze position. This firstly validated our artifact detection: gaze deviated very little from central fixation (<0.1°). Second, we did not observe systematic gaze biases toward the position of either the selected or non-selected shapes during the delay period. This rules out miniature gaze patterns as a confound to our decoding findings. Colors indicate the position bins used in the inverted encoding modeling procedures. Error bars reflect standard error of the mean.

